# *Trp53* ablation fails to prevent microcephaly in mouse pallium with impaired minor intron splicing

**DOI:** 10.1101/2021.03.05.434172

**Authors:** Alisa K. White, Marybeth Baumgartner, Madisen F. Lee, Kyle D. Drake, Gabriela S. Aquino, Rahul N. Kanadia

## Abstract

Mutations in minor spliceosome component RNU4ATAC, a small nuclear RNA (snRNA), are linked to primary microcephaly. We have reported that in the conditional knockout (cKO) mice for *Rnu11*, another minor spliceosome snRNA, minor intron splicing defect in minor intron-containing genes (MIGs) regulating cell cycle resulted in cell cycle defects, with a concomitant increase in γH2aX+ cells and p53-mediated apoptosis. *Trp53* ablation in the *Rnu11* cKO mice did not prevent microcephaly. However, RNAseq analysis of the double knockout (dKO) pallium reflected transcriptomic shift towards the control from the Rnu11 cKO. We found elevated minor intron retention and alternative splicing across minor introns in the dKO. Disruption of these MIGs resulted in cell cycle defects that were more severe and detected earlier in the dKO, but with delayed detection of γH2aX+ DNA damage. Thus, p53 might also play a role in causing DNA damage in the developing pallium. In all, our findings further refine our understanding of the role of the minor spliceosome in cortical development and identify MIGs underpinning microcephaly in minor spliceosome-related diseases.

## Introduction

The complex higher-order functions such as controlling motor and sensory functions as well as perception, thought, and memory carried out by the cortex depends on a complex neuronal architecture (Douglas et al., 1995). The cortex consists of a vast number of diverse neuronal subtypes that must be in the right proportion that are laminated with the appropriate synaptic partners. Thus, proper cortical development is crucial for proper cortical function. Proper early brain development requires a precise balance of progenitor cell amplification and neurogenesis, which places cell cycle regulation at the core of cortical development and disease (Dehay and Kennedy, 2007). Indeed, a range of sources, including environmental factors including maternal exposure to toxins and pathogens have been linked to disruption in cell cycle and cortical development (Chen et al., 2003; Feldman et al., 2012; von der Hagen et al., 2014; Rasmussen et al., 2016). Consequently, proliferation defect and too few progenitor cells often manifests as primary microcephaly in humans.

Mutations in a gene(s) involved in cell cycle that result in microcephaly in humans provide insight into the cellular and molecular processes essential for cortical development (von der Hagen et al., 2014). Amongst the genes linked to microcephaly, more than half encode proteins localizing to centrosome and are vital for centriole biogenesis and duplication (Jayaraman et al., 2018). Disruptions in such genes would result in centrosome dysfunction, impairing chromosome segregation. Furthermore, many genes involved in DNA replication have been associated with microcephaly through replicative stress, leading to DNA damage (Jayaraman et al., 2018). Moreover, genes regulating cell cycle checkpoints, such as the spindle assembly checkpoint, have also been linked to primary microcephaly, highlighting the importance of cell cycle regulation for proper cortical development.

Rare forms of primary microcephaly are linked mutations in *RNU4ATAC*, which encodes the U4atac small nuclear RNA (snRNA), an essential component of the minor spliceosome. This disruption results in a spectrum of diseases including microcephalic osteodysplastic primordial dwarfism type 1 (MOPD1), Lowry-Wood syndrome (LWS), and Roifman syndrome (RS) (Edery et al., 2011; He et al., 2011; Merico et al., 2015; Farach et al., 2018). Despite the variability in the severity of these diseases, microcephaly is common to all, which reveals the significance of the minor spliceosome in cortical development. The minor spliceosome, consisting of U11, U12, U4atac, U6atac, and U5 snRNAs and associated proteins is responsible for the removal of less than 0.5% of all introns, termed minor introns, found in 650 genes in mouse(Levine and Durbin, 2001; Patel and Steitz, 2003; Olthof et al., 2019). Minor intron-containing genes (MIGs) perform disparate functions, ranging from transcription factors and splicing factors to protein kinases, translation regulators, ion channels, and others (Burge et al., 1998; Baumgartner et al., 2019). As such, disruption of minor intron splicing would result in a large molecular footprint and is therefore difficult to determine the precise molecular defect that drives microcephaly in these patients. We have previously leveraged our conditional knockout (cKO) mouse targeting *Rnu11*, the gene encoding the U11 snRNA to explore the molecular and cellular defects underpinning cortical defects in these diseases (Baumgartner et al., 2018). We reported that *Emx1*-Cre-mediated ablation of *Rnu11* resulted in microcephaly at birth, which was caused by simultaneous cell cycle defects and cell death of radial glial cells (RGCs) in the U11-null developing cortex (pallium) (Baumgartner et al., 2018).

Here we test the hypothesis that the primary defect driving microcephaly in the U11-null developing cortex is cell cycle defect, which is upstream of the cell death phenotype. Several mouse models of microcephaly, including *Sas4* (Bazzi and Anderson, 2014; Insolera et al., 2014), *Kif20b* (Janisch et al., 2013; Little and Dwyer, 2019), *Magoh (Silver et al., 2010; Mao et al., 2016)*, display cell cycle defects and p53-mediated cell death. Microcephaly in these mouse models can be rescued through ablation of p53. Given that we also observed cell cycle defects and p53-mediated apoptosis in the U11-null pallium, we sought to prevent cell death by ablating *Trp53* (p53) in the *Rnu11* cKO mouse to generate double knockout (dKO) mice. This allowed us to block apoptosis in the U11-null pallium and study both the progression of DNA damage and cell cycle defects triggered by minor spliceosome disruption and the relative contribution of DNA damage and cell cycle defects on microcephaly.

Here we report that blocking cell death failed to prevent microcephaly in the dKO mice. This finding suggests that microcephaly in minor spliceosome diseases is primarily driven by cell cycle defects, which were detected earlier and more severe in the dKO compared to the *Rnu11* cKO. Specifically, RGCs in the dKO showed prometaphase elongation, decrease in the number of S-phase RGCs, and a significant increase in overall cell cycle length, driven primarily by lengthening of G1 and/or G2 phases. Importantly, we identified the primary molecular defect of elevated minor intron retention in the dKO, which was similar to that in the *Rnu11* cKO. We also observed elevated alternative splicing (AS) across minor introns in both the *Rnu11* cKO and dKO developing cortices, which is consistent with our recent report (Olthof et al., 2021). Based on current literature and previous observations in the *Rnu11* cKO pallium, we anticipated that accumulation of DNA damage, which is thought to activate p53 stabilization, would be detected in the dKO (Lakin and Jackson, 1999; Cheng and Chen, 2010). Surprisingly, DNA damage was significantly delayed in the dKO, relative to the *Rnu11* cKO. This finding suggests that in the developing U11-null pallium, p53 might not only respond to DNA damage, but might also participate in DNA damage. In all, here we uncovered that microcephaly in minor spliceosome disease is driven primarily by cell cycle defects, predominantly those affecting mitotic progression.

## Results

### Ablation of p53 in Rnu11 cKO does not prevent microcephaly

Shown in Figure 1A is a model summarizing the molecular and cellular defects in the *Rnu11* cKO mouse that result in primary microcephaly (Fig. 1A)(Baumgartner et al., 2018). To block cell death by ablating P53, we produced mouse-lines with the following genotypes: *Rnu11*^WT/Flx^::Trp53^WT/WT^::*Emx1*-Cre^+^ (control), *Rnu11*^Flx/Flx^::*Trp53* ^WT/WT^ ::*Emx1*-Cre^+^ (*Rnu11* cKO), and *Rnu11*^Flx/Flx^::*Trp53* ^Flx/Flx^ ::*Emx1*-Cre^+^ (dKO) mice (Fig. 1B). At birth, we observed no rescue of microcephaly in the dKO mice (Fig. 1C), which was confirmed by cortical weight measurements (Fig. 1C’). To confirm this we performed quantification of Nissl-stained sections across the rostral to caudal axis in control, *Rnu11* cKO, and dKO brains which revealed significantly reduced mediolateral neocortical length in the *Rnu11* cKO and dKO compared to the control. However, comparison of the *Rnu11* cKO and dKO cortices revealed a slight, but significant increase in the mediolateral neocortical length and medial neocortical thickness in the dKO, indicative of partial rescue of microcephaly (Fig. 1D’-D’’).

**Figure 1.**
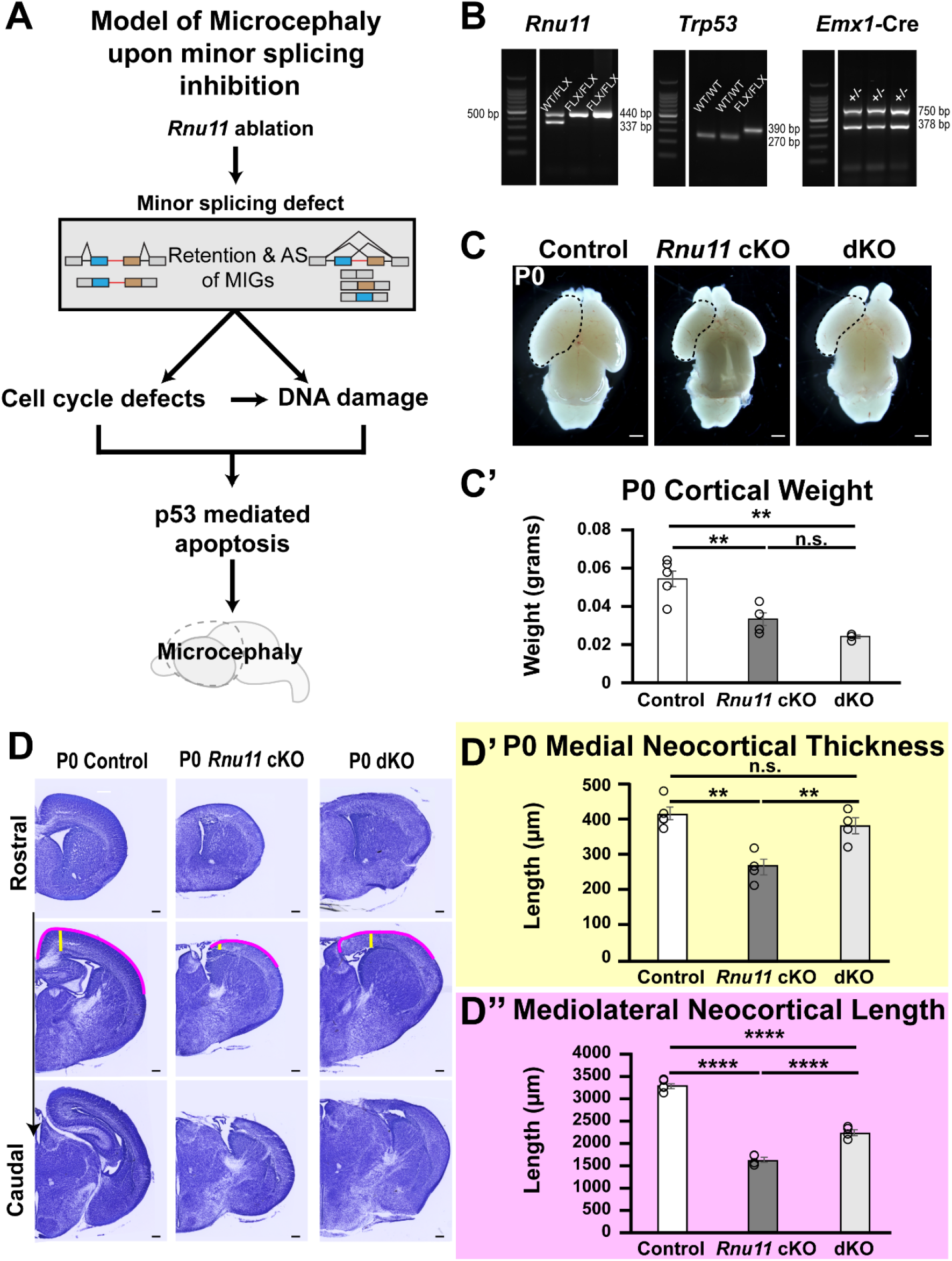
Removal of *Trp53* in the *Rnu11* cKO mouse does not fully rescue microcephaly. **(a)** Schematic of the impacts of U11 loss in the *Rnu11* cKO mouse pallium, in which the mis-splicing of MIGs leads to cell cycle defects, DNA damage, and p53-mediated cell death, ultimately resulting in microcephaly. Model informed by Baumgartner et al. 2018(Baumgartner et al., 2018). **(b)** Agarose gels for genotype verification of control, *Rnu11* cKO, and dKO mice (left to right within each gel) for *Rnu11* (left), *Trp53* (middle), and *Emx1*-cre (right). Ladder 100 bp MW (Promega). **(c)** Images of postnatal day 0 (P0) brains of control (left), *Rnu11* cKO (middle), and dKO (right) with left cortices traced in black dotted lines. Scale bar=1mm. **(d)** Representative bright field images of Nissl-stained 50µm thick cryosections of control (left), *Rnu11* cKO (middle), and dKO (right) brains from rostral (top) to caudal (bottom). Scale bar=200µm. Yellow and pink lines represent medial neocortical thickness and mediolateral neocortical length quantification strategies, respectively. **(d’-d’’):** Bar graphs of **(d’)** medial neocortical thickness and **(d’’)** mediolateral neocortical length at P0 of control (white, left), *Rnu11* cKO (black, middle), and dKO (grey, right). Data are presented as mean±s.e.m. Statistical significance determined by one-way ANOVA followed by post-hoc Tukey test. ****=*P*<0.0001, ***=*P*<0.001, **=*P*<0.01, n.s.=not significant.

### p53 ablation in Rnu11 cKO drives transcriptome towards control

In the *Rnu11* cKO pallium, stabilization of p53 protein resulted in detection of p53 by E12 (Baumgartner et al., 2018). To verify p53 loss in the dKO pallium, we performed IHC for p53 on E14.5 pallial sections. While p53 protein was detected in the E14.5 *Rnu11* cKO pallium, we did not observe p53 signal in the dKO (Figure 2A). Since p53-mediated apoptosis was the sole GO Term enriched for by the upregulated genes in the *Rnu11* cKO as previously reported, we explored the impact of p53 ablation on the overall transcriptome. We have previously reported that cell death in the *Rnu11* cKO pallium begins at E12 and reaches its peak at E14. Therefore, we hypothesized that blocking cell death in the dKO would have the strongest molecular impact at E13.5 and performed bulk Ribo-Zero total RNAseq of the pallium at E13.5 (n=4, per genotype). Confirmation of clean pallial dissection was determined through expression of known markers of the region. Expression of *Emx1* and another known dorsal telencephalon marker, *Lhx2*, was high, whereas non-dorsal telencephalon markers from neighboring regions, such as *Dbx1* and *Lhx6*, were not expressed (Table S1)(Grant et al., 2012). Next, we confirmed the genetic manipulation by identifying downregulation of *Rnu11* in the *Rnu11* cKO and downregulation of both *Rnu11* and *Trp53* in the dKO, relative to the control (Table S1). The level of reduction of both *Rnu11* and *Trp53* is commensurate with the number of *Emx1*-cre negative cells captured in our dissection (Bartolini et al., 2013; Baumgartner et al., 2018).

**Figure 2.**
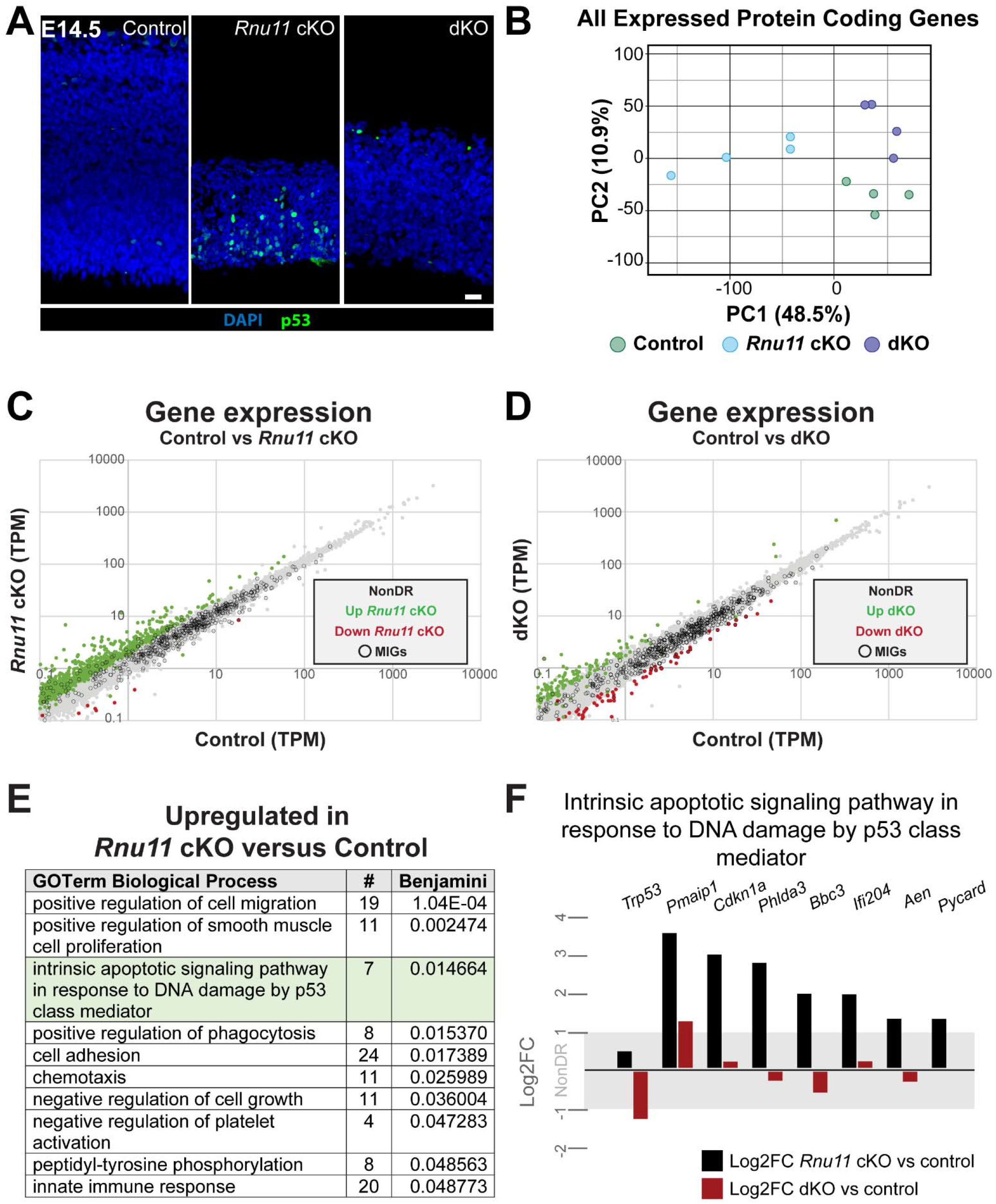
Removal of p53 rescues the pro-apoptotic molecular phenotype observed in the *Rnu11* cKO pallium. **(a)** Immunohistochemistry (IHC) for p53 (green) in E14.5 sagittal sections of the control (left), *Rnu11* cKO (middle), and dKO (right) pallium. Nuclei are marked by DAPI (blue). Scale bar=20µm. **(b)** Principal component analysis (PCA) of all expressed (transcripts per million [TPM]≥1) protein-coding genes in control (green circles, n=4), *Rnu11* cKO (blue circles, n=4), and dKO (purple circles, n=4). **(c)** Scatterplot of gene expression (TPM) of protein-coding genes in the control (x-axis) and *Rnu11* cKO (y-axis) in the E13.5 pallium. Genes non-differentially regulated (NonDR) in grey, upregulated in the *Rnu11* cKO in green, downregulated in the *Rnu11* cKO in red, and MIGs outlined with black circles. **(d)** Scatterplot of gene expression (TPM) of protein-coding genes in the control (x-axis) and dKO (y-axis) in E13.5 pallium. Genes non-differentially regulated (NonDR) in grey, upregulated in the dKO in green, downregulated in the dKO in red, and MIGs outlined with black circles. **(e)** Table of GO Terms with significant enrichment by genes upregulated in the E13.5 *Rnu11* cKO compared to the control, identified by DAVID analysis. The term “intrinsic apoptotic signaling pathway in response to DNA damage by p53 class mediator” is highlighted in green. **(f)** Bar chart of Log2 fold-change (FC) values of the 7 genes enriching for the green highlighted GO Term in the *Rnu11* cKO vs control comparison (black), and their Log2FC in the dKO vs control comparison (red). The gray horizontal bar indicates the range of non-differential expression (NonDR) calls.

Principal component analysis (PCA) on the three genotypes based on expression of all protein-coding genes revealed that the dKO and control transcriptomes were more similar than that of the *Rnu11* cKO (PC1 48.5%, Fig. 2B). This suggested that activation of p53, a transcription factor, was contributing to the transcriptomic signature of the *Rnu11* cKO, likely driven by the activation of pro-apoptotic pathways. To identify precise transcriptomic shifts in the *Rnu11* cKO pallium, we performed differential gene expression analysis in the control compared to the *Rnu11* cKO (Fig. 2C), which identified a large set of genes (401) that were upregulated in the *Rnu11* cKO. However, fewer genes were upregulated in the dKO compared to the control (Fig. 2C&D). In contrast, more genes (32, including 13 MIGs) were downregulated in the dKO, compared to the 3 downregulated genes, including 1 MIG (*Lage3*), in the *Rnu11* cKO (Fig. 2C&D). The downregulated genes in the dKO enriched for more broad terms including nucleosome, chromosome, chromatin, and protein heterodimerization activity, which is consistent with the PCA results (Fig. 2B).

Database for Annotation, Visualization, and Integrated Discovery (DAVID) (Huang da et al., 2009) analysis of the 401 genes upregulated in the E13.5 *Rnu11* cKO pallium significantly enriched for the GO Term “intrinsic apoptotic pathway through p53 class mediator in response to DNA damage” (Fig. 2E). The genes underpinning this enrichment included *Pmaip1, Phlda3, Bbc3*, and *Aen*, which are downstream targets of p53 in the apoptotic pathway (Fig. 2F). In contrast, similar assessment of the 43 genes upregulated in the dKO compared to the control did not reveal any significant functional enrichments. We hypothesized that the p53 effector genes upregulated in the *Rnu11* cKO would be rescued in the dKO. We plotted Log2Fold Change (FC) of the 7 p53-effector genes in both the *Rnu11* cKO and dKO compared to the control (Fig. 2F). All genes except *Pmaip* were non-differentially expressed in the dKO when compared to the control (Fig. 2F, red bars, Table S1).

### Minor intron splicing defect is not rescued by p53 ablation

Ablation of p53 blocked the transcriptional upregulation of p53-mediated apoptotic effectors, yet it did not prevent microcephaly in the dKO (Fig. 1). Therefore, we next sought to determine whether the expected primary defect, impaired minor splicing, was present in the dKO. As anticipated, both the *Rnu11* cKO and dKO displayed significantly elevated minor intron retention, quantified by mis-splicing index (MSI), relative to the control (Fig. 3A,(Baumgartner et al., 2018)). In contrast, the median MSI of minor introns was not significantly different between the *Rnu11* cKO and dKO (Fig. 3A). Notably, PCA on minor intron retention (%MSI) values revealed clustering of the *Rnu11* cKO and dKO samples, indicating that the pattern of minor splicing defects was not affected by p53 ablation (Fig. 3B). Specifically, we identified 265 MIGs with significantly increased minor intron retention in the *Rnu11* cKO compared to the control, and 231 MIGs with significantly increased minor intron retention in the dKO compared to the control (Fig. S1A). There were 210 events shared between the *Rnu11* cKO and the dKO compared to the control (Fig. S1A). DAVID analysis on MIGs with elevated minor intron retention enriched for multiple GO Terms, including cell cycle, cellular response to DNA damage stimulus, and mitotic nuclear division (Fig. S1A). DAVID analysis on the *Rnu11* cKO-specific and the dKO-specific MIGs with elevated minor intron retention independently revealed general terms for the *Rnu11* cKO-specific MIGs, while no GO Terms were significantly enriched for the MIGs in the dKO (Fig. S1A).

**Figure 3.**
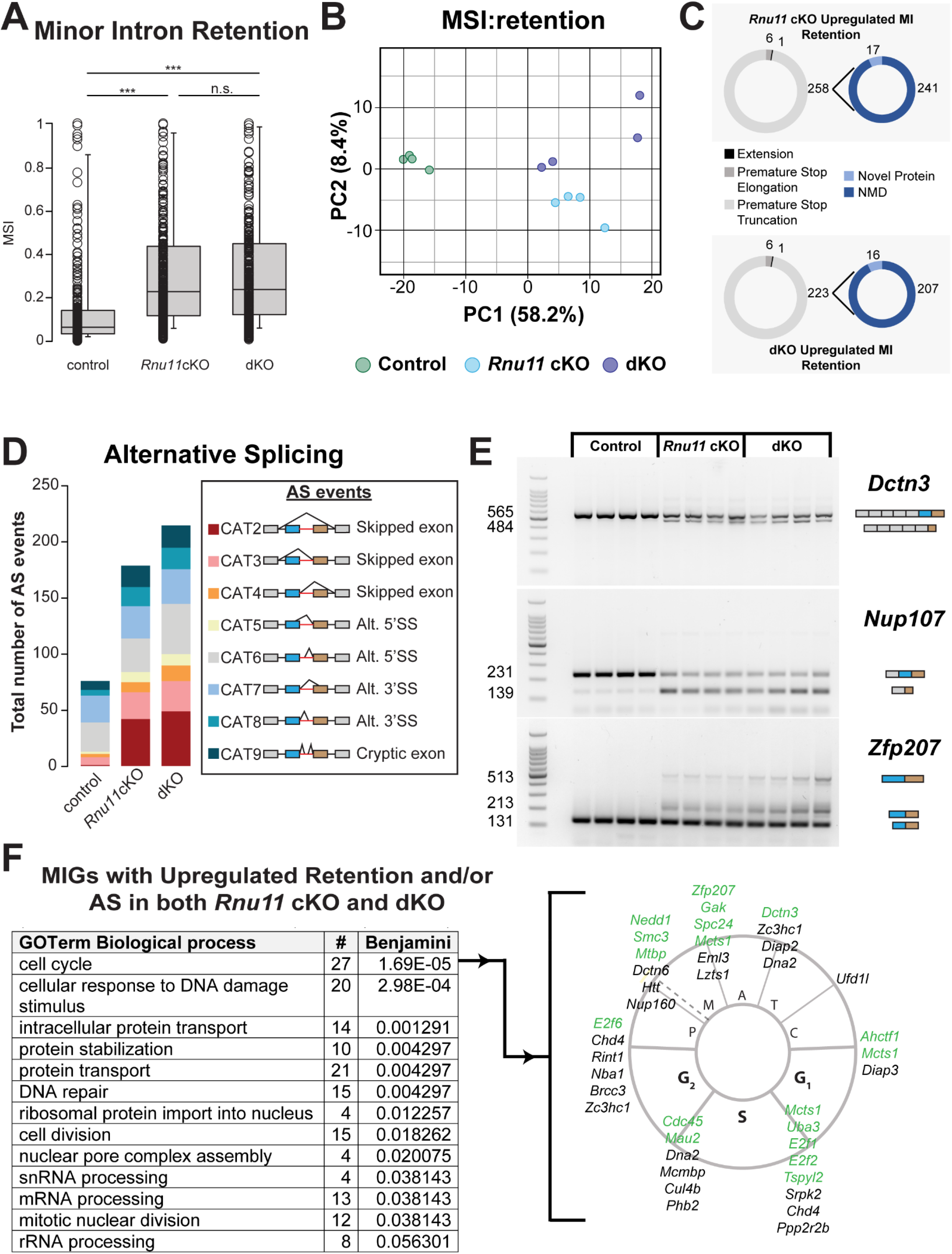
Primary defect of mis-splicing of minor intron containing genes involved in cell cycle is comparable in the *Rnu11* cKO and dKO. **(a)** Boxplot showing the 10^th^ to 90^th^ percentiles of the median mis-splicing index (MSI) for all (402) minor introns that show retention in the control, *Rnu11* cKO and dKO. Statistical significance was determined by Kruskal-Wallis H test. ***=*P*<0.001, n.s.=not significant. **(b)** PCA of minor intron retention (minor intron MSI) in control (blue circles, n=4), *Rnu11* cKO (green circles, n=4), and dKO (purple circles, n=4). **(c)** At left, pie charts of predicted consequences of minor intron retention on the open reading frame (ORF) of MIGs with elevated minor intron retention in the *Rnu11* cKO (top left, greyscale) and the dKO (bottom left, greyscale) relative to control. At right, pie charts representing whether premature stop codons due to minor intron retention are predicted to trigger NMD or novel protein production in the *Rnu11* cKO (top right, blue) and dKO (bottom right, blue). **(d)** Bar graph of alternative splicing (AS) events around minor introns in the control, *Rnu11* cKO, and dKO. Schematics of the different types of AS events color-coded in the bar graph are displayed in the key on the right. **(e)** Agarose gel images of selected AS events around the minor intron for cell cycle-related MIGs as detected by reverse transcriptase-PCR (RT-PCR) (*Dctn3* top, *Nup107* middle, *Zfp207* bottom). **(f)** Table of Biological Process GO Terms enriched by MIGs with shared upregulation of minor intron retention and/or alternative splicing in both the *Rnu11* cKO and dKO compared to the control. In the cell cycle schematic at right, MIGs enriching for the top biological process (cell cycle) are listed in green text and located within the specific cell cycle stage(s)/transition(s) they associate. MIGs listed in black text also regulate cell cycle, do not enrich for the specific GO Term, and are similarly distributed in the cell cycle schematic.

Often, retention of minor introns is predicted to introduce a premature stop codon, resulting in nonsense-mediated decay (NMD), nuclear degradation, or the production of aberrant protein (Nagy and Maquat, 1998; Houseley and Tollervey, 2009; Olthof et al., 2021). To further parse the functional impacts of elevated minor intron retention on each MIG, we analyzed the open reading frame (ORF) of all MIGs with significantly elevated minor intron retention in at least one of the two conditions (Baumgartner et al., 2018). Of the 265 MIGs with increased minor intron retention in the *Rnu11* cKO compared to the control, the majority (258/265; 97.4%) of the retained transcripts were predicted to contain a premature stop codon (Fig. 3C). Of those with premature stop codons, 241 were predicted to be targeted for degradation via NMD, while 17 of these transcripts were predicted to be translated and produce truncated proteins (Fig. 3C). Similarly, of the 231 MIGs with significantly elevated minor intron retention in the dKO relative to the control, the majority (223/230, 97.0%) were predicted to introduce a premature stop codon. Of those predicted to contain a premature stop codon, 207 were predicted to be targets for NMD, while the remaining 16 were predicted to result in the production of truncated proteins (Fig. 3C).

The potential targeting of MIG transcripts with minor intron retention by NMD would prevent protein production from these transcripts, thus driving severe repression of the cellular functions performed by these MIGs. To identify these MIG-regulated functions, we performed GO Term enrichment analysis using the shared and unique lists of MIGs with significantly elevated minor intron retention predicted to be targets of NMD. Similar to our analysis of all MIGs with elevated minor intron retention, we found that the MIGs with elevated minor intron retention in both the *Rnu11* cKO and dKO significantly enriched for terms including cell cycle, cell division, and cellular response to DNA damage stimulus (Fig. S1B). As before, the MIGs with unique events in the *Rnu11* cKO and dKO enriched only for protein transport and mRNA transport, or had no significant enrichments, respectively (Fig. S1B). This suggested that the underlying molecular signature shared in the *Rnu11* cKO and dKO was unaffected upon p53 ablation.

In addition to minor intron retention, disruption of the minor spliceosome can trigger major spliceosome-mediated alternative splicing (AS) around minor introns (Olthof et al., 2021). Utilizing a previously published, minor intron-focused alternative splicing-specific MSI calculation strategy, we identified 76, 179, and 215 AS events in the control, *Rnu11* cKO, and dKO, respectively. Of these AS events, 119 were upregulated in the *Rnu11* cKO compared to the control, and 156 in the dKO compared to the control (Fig. 3D; Fig. S1C). In both the *Rnu11* cKO and dKO this increase in AS events was primarily observed through exons skipping events (Fig. 3D, CAT2, 3, 4). Notably, we observed the greatest increase in the skipping of the upstream and downstream exons flanking the minor intron (CAT2), which was detected only once in the control, however, 42 events were detected in the *Rnu11* cKO and 49 in the dKO. Moreover, there were few AS events, including (CAT6, 7) that occurred at high levels in the control (26, 24) and were further increased in both the *Rnu11* cKO (30, 29) and dKO (45, 31).

Of these upregulated events in the *Rnu11* cKO and dKO relative to the controls, there were 105 shared AS events. This upregulation of AS around the minor intron was confirmed and visualized for select MIGs through reverse transcriptase-PCR (RT-PCR) analysis (Fig. 3E). For instance, the canonically spliced transcript of *Dctn3* was detected across all conditions (Fig. 1E, 565bp), however, an isoform that skips both exons flanking the minor intron was only observed in the *Rnu11* cKO and the dKO samples (Fig. 3E; CAT3, 484bp) (Olthof et al., 2021). In case of *Nup107*, an additional isoform (139bp) to the canonically spliced product (231bp) was detected in the *Rnu11* cKO and dKO (Fig. 3E; CAT3 + CAT7). This AS event was the result of skipping of the exon upstream of the minor intron and the use of an alternative 3’ splice site in the exon downstream of the minor intron which was observed in both the *Rnu11* cKO and dKO (Olthof et al., 2021). Finally, *Zfp207* in addition to the presence of the canonical splicing event around the minor intron (131bp), we validated two AS events around the minor intron, one utilizing alternative 5’ and 3’ splice site usage (Fig. 3E; CAT6 + CAT8, 513bp) and another with only alternative 5’ splice usage (CAT6, 213bp). GO Term analysis of the MIGs with shared, upregulated AS events in the *Rnu11* cKO and dKO significantly enriched for terms including nucleus, kinetochore, and condensed chromosome kinetochore (Fig. S1C).

Next, we combined the lists of MIGs with shared, upregulated AS events around minor introns with the list of shared MIGs displaying elevated minor intron retention in both the *Rnu11* cKO and dKO. This combined list of mis-spliced MIGs was then subjected to DAVID analysis. The most significantly enriched GO Term by the shared mis-spliced MIGs was cell cycle (Fig. 3F). The MIGs underlying this enrichment were found to be involved in all stages of the cell cycle (Fig. 3F, green text). Notably, we found several MIGs that were missed by DAVID enrichment that had literature support for their involvement in regulating various stages of the cell cycle (Fig. 3F, black text).

### Mitotic progression of RGCs is exacerbated in the dKO

The shared splicing defects in both the *Rnu11* cKO and dKO pallium predicted the disruption of cell cycle. Moreover, functional enrichment analyses suggested that mitosis would be particularly affected in these mice. In our previous report, we found that the RGC population displayed the most significant cell cycle defects, p53 protein upregulation, and cell death in the U11-null pallium (Baumgartner et al., 2018). To assess mitosis in U11-null RGCs, we performed IHC for the mitotic marker Aurora B and Pax6, to mark RGCs (Gotz et al., 1998), in E12.5, E13.5 and E14.5 pallia from control, *Rnu11* cKO, and dKO embryos. For mitotic stage calls, we then utilized the staining pattern of Aurora B to quantify the percent of mitotic RGCs in each phase of mitosis (Sun et al., 2008; Baumgartner et al., 2018). At E12.5, there was a significant increase in the percent of mitotic RGCs in prometaphase in both the *Rnu11* cKO and the dKO compared to the control (Fig. 4A&B). Additionally, there was a significantly lower fraction of mitotic RGCs in metaphase in the dKO when compared to both the *Rnu11* cKO and the control (Fig. 4B). At this timepoint (E12.5) the percent of RGCs in prophase, anaphase, and telophase were unaffected across all genotypes. By E13.5, there was a further increase in the percentage of mitotic RGCs in prometaphase in both the *Rnu11* cKO and the dKO, relative to the control, which was concomitant with a significant reduction in the percentage of mitotic RGCs in metaphase (Fig. 4A&B). In the E13.5 dKO, we also observed a significant decrease in the proportion of mitotic RGCs in telophase, compared to the *Rnu11* cKO and control. At this timepoint (E13.5), the percentage of mitotic RGCs in prophase and anaphase was not significantly impacted in the *Rnu11* cKO and dKO compared to the control. When we extended this analysis to E14.5, we observed continued increase in the proportion of mitotic RGCs in prometaphase in both the *Rnu11* cKO and the dKO relative to the control, which coincided with significant reductions in the percentage of mitotic RGCs in both metaphase and telophase. At this timepoint (E14.5), the fraction of mitotic RGCs in prophase and anaphase were still comparable across all genotypes.

**Figure 4.**
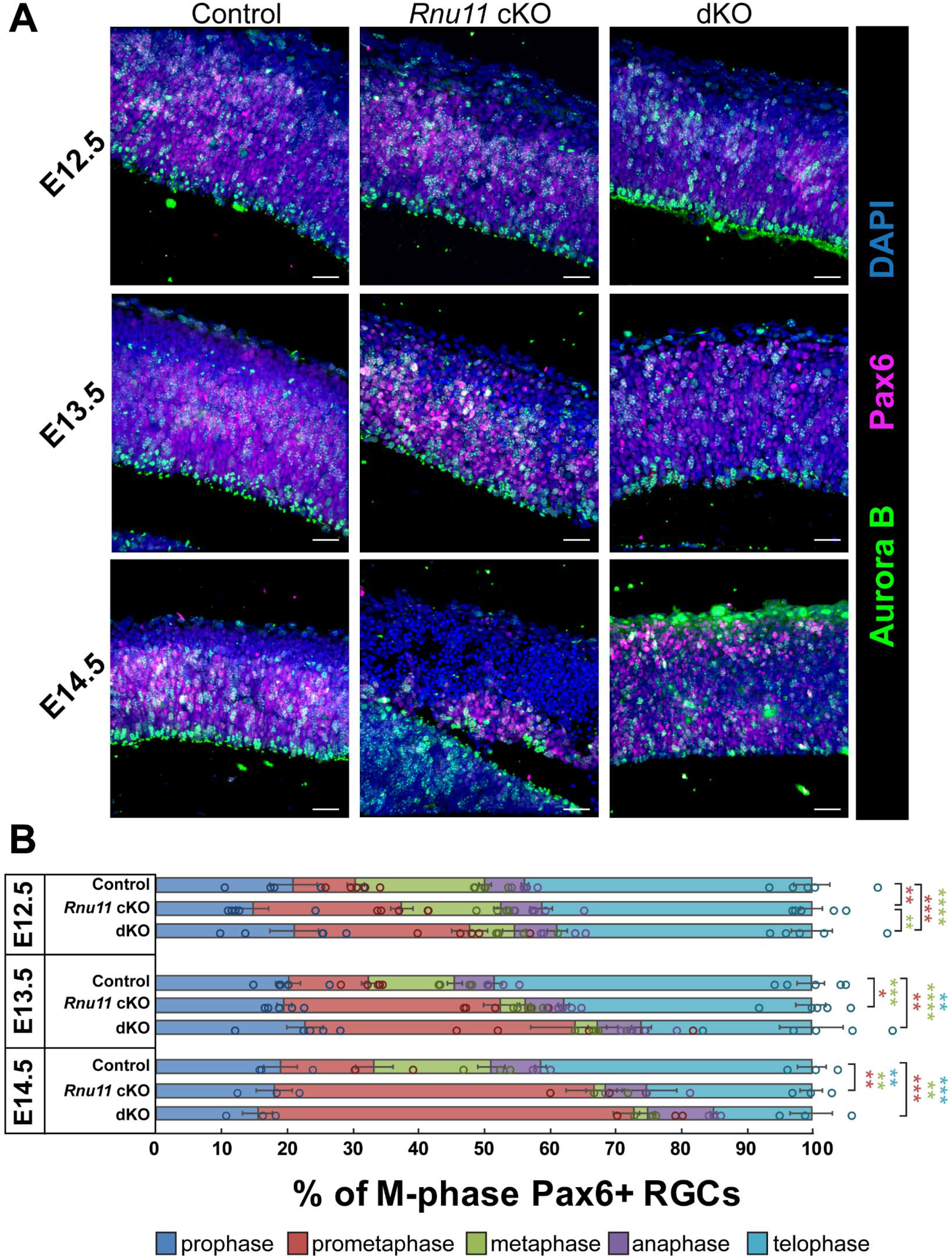
Mitotic progression of radial glial cells (RGCs) is significantly impacted in both *Rnu11* cKO and dKO by E12.5. **(a)** Immunohistochemistry (IHC) for Aurora B (green) and Pax6 (magenta) on sagittal sections of control (left), *Rnu11* cKO (middle), and dKO (right) pallium across E12.5 (top), E13.5 (middle), E14.5 (bottom). Nuclei are marked by DAPI (blue). Scale bars=30μm. **(b)** Quantification of the percentage of Aurora B+ mitotic RGCs (Pax6+) across different mitotic phases in control, *Rnu11* cKO, and dKO across E12.5, E13.5, and E14.5. Data are presented as mean±s.e.m. Statistical significance was determined by one-way ANOVA, followed by post-hoc Tukey test.**P<0.01; ***=*P*<0.001. Asterisks color indicates the specific comparison tested for statistical significance, corresponding to the bar color-coding scheme.

### Cell cycle length is increased in dKO

The U11-null cells in the *Rnu11* cKO and dKO particularly experience defects in prometaphase-to-metaphase progression (Fig. 4). However, MIGs involved in S-phase are also mis-spliced in both the *Rnu11* cKO and the dKO pallium (Fig. 3F). Since many of S phase-linked MIGs function in DNA replication, disruption of these MIGs could impact the speed and accuracy of DNA replication. Thus, we sought to interrogate RGCs in S-phase, as well as determine S-phase and total cell cycle length. To do this we utilized a strategy in which we pulsed pregnant dams once with BrdU 2 hours before harvest, then pulsed once with EdU 30 minutes prior to harvest (Fig. 5A)(Martynoga et al., 2005). The percent of RGCs in S-phase upon harvest was determined by the percent of Pax6+ RGCs that were also EdU+ out of all Pax6+ cells. At E12.5, the percent of RGCs in S-phase was comparable across all genotypes (Fig. S2). However, there was a significant decrease in the percentage of S-phase RGCs in the dKO compared to the control by E13.5. By E14.5, there was a further reduction in the percentage of RGCs in S-phase in both the *Rnu11* cKO and the dKO relative to the control (Fig. S2).

**Figure 5.**
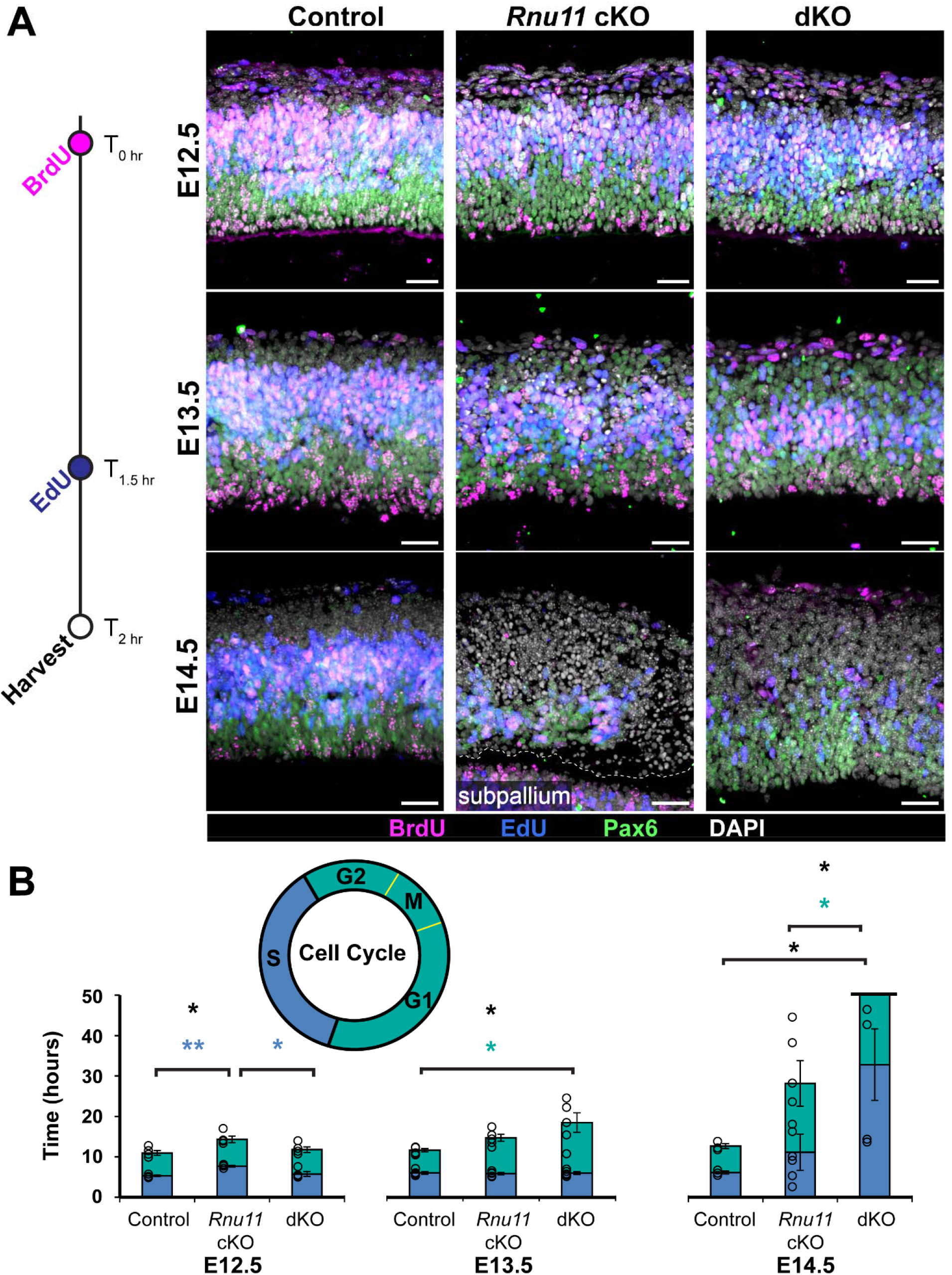
Total cell cycle length of RGCs is significantly extended in the dKO by E13.5 and E14.5, respectively. **(a)** BrdU/EdU injection paradigm adapted from Martynoga et al. to quantify total cell cycle speed and S-phase length of RGCs (Martynoga et al., 2005). A single BrdU injection (magenta) was performed at T=0 hours, followed by a single EdU injection (blue) at T=1.5 hours, with collection at T=2 hours (white). Subsequent immunohistochemistry (IHC) detection for BrdU and Pax6 (green) were performed in conjunction with EdU detection in sagittal pallial sections of the control (left), *Rnu11* cKO (middle), and dKO (right) at E12.5 (top), E13.5 (middle), and E14.5 (bottom). Nuclei are stained by DAPI (white). Scale bars=30μm. **(b)** Quantified S-phase length (blue), G2-G1 length (turquoise) and total cell cycle (full bar, outlined in black) length in the control, *Rnu11* cKO, and dKO pallium across E12.5, E13.5, and E14.5, calculated using the Martynoga et al. method (Martynoga et al., 2005). Data are presented as mean± s.e.m. Statistical significance was determined by one-way ANOVA, followed by post-hoc Tukey test. n.s., not significant, *=*P*<0.05; **=*P*<0.01; ***=*P*<0.001. Asterisks color indicates the specific comparison tested for statistical significance, corresponding to the bar color-coding scheme.

To interpret these shifts in S-phase RGC population, we calculated S-phase and the total cell cycle length in RGCs, by utilizing both pulses of BrdU and EdU (Fig. 5A). This was followed by IHC for Pax6 and BrdU, coupled with EdU detection. We calculated estimated lengths of S-phase and total cell cycle in RGCs in accordance with Martgyar et al.(Martynoga et al., 2005). At E12.5, there was a significant increase in the calculated length of S-phase as well as total cell cycle length in the *Rnu11* cKO compared to the control (Fig. 5B, blue fill and black line). By E13.5, the significant increase in S-phase and cell cycle length was no longer observed in the *Rnu11* cKO. However, in the dKO there was a significant increase in the length of total cell cycle and in the length of G2-G1 compared to the control (Fig. 5B). Finally, at E14.5, total cell cycle length was significantly increased in the dKO compared to both the control and the *Rnu11* cKO (Fig. 5B). Calculations of estimated G2-G1 and total cell cycle length in the dKO were too high for them to be physiologically feasible (>30 hours), indicating that RGCs most likely had arrested in S-phase in the dKO and, more dramatically, G2-G1 phases (Fig. 5B). Delay in G2-G1 phases was consistent with the significant decrease in the S-phase RGC population at this time-point (Fig. S2). In contrast, the lengths of S-phase, G2-G1 phases, and the total cell cycle in the *Rnu11* cKO were not significantly different from the control at E14.5.

### Removal of Trp53 in Rnu11 cKO, prevents cell death and delays DNA damage accumulation

Ablation of p53 was performed to block cell death in response to DNA damage and cell cycle defects in the *Rnu11* cKO pallium. Therefore, we next investigated cell death and DNA damage accumulation in the dKO. For this, we performed terminal deoxynucleotidyl transferase dUTP nick-end labeling (TUNEL), a late-stage cell death marker (Kyrylkova et al., 2012), coupled with IHC for Cleaved-Caspase 3 (CC3) (Faleiro et al., 1997), a marker of early cell death, and γH2AX to detect DNA damage (Sharma et al., 2012). We observed that ablation of p53 successfully blocked cell death, as we observed no difference in TUNEL+ cells or CC3+ cells in the dKO at E12.5, E13.5, and E14.5 compared to the control (Fig. 6A&B). As expected, we observed significant elevation of death, using both markers, in the *Rnu11* cKO compared to both the control and dKO at all time-points (Fig. 6A&B). Unexpectedly, DNA damage, generally thought to drive p53 protein stabilization and p53-mediated apoptosis, was delayed in the dKO, such that significant DNA damage marked by γH2AX was not detected until E14.5 in the dKO compared to the control, which was further increased by E15.5 (Fig. 6A&C, Fig. S3).

**Figure 6.**
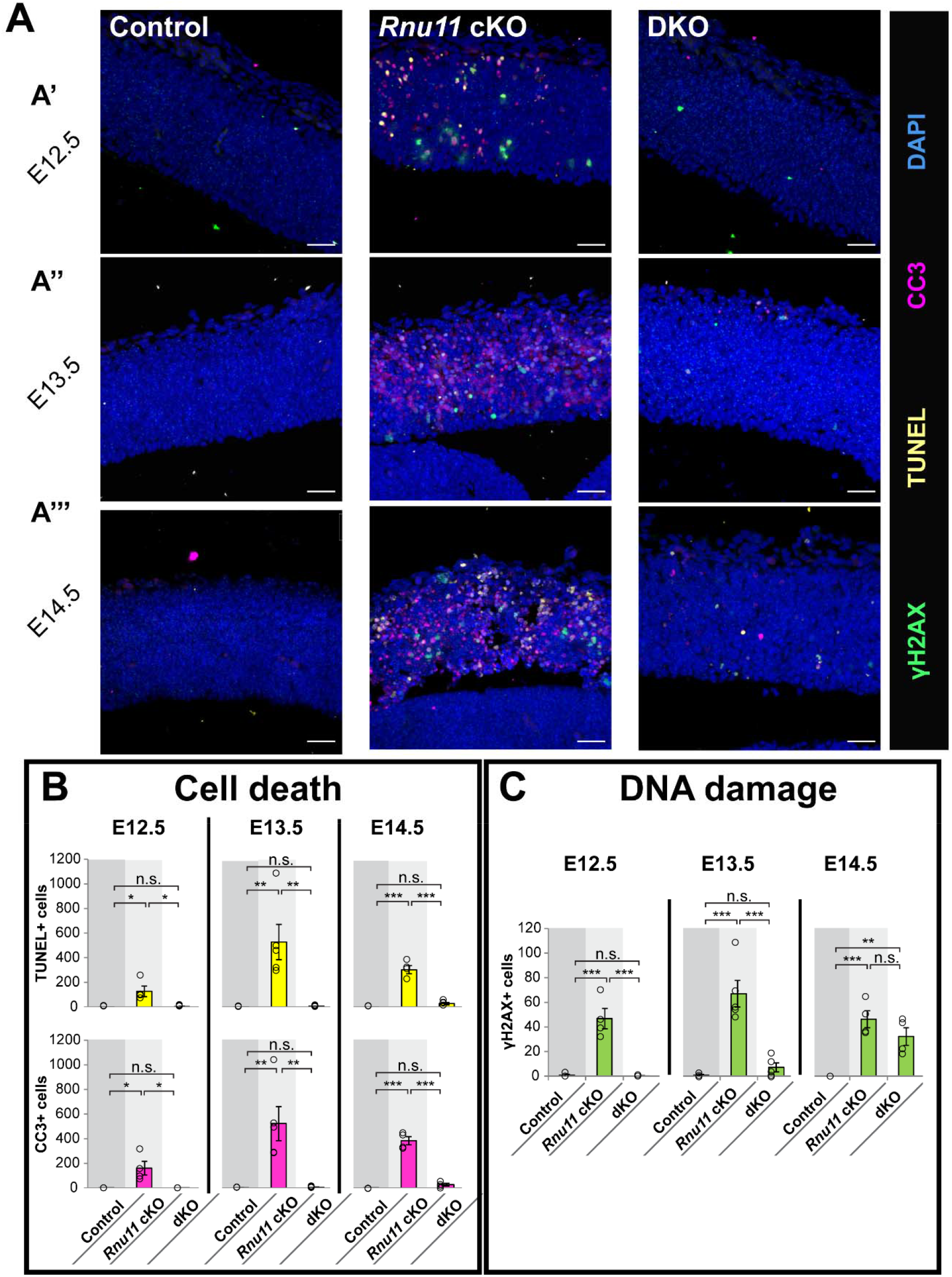
Removal of p53 in the *Rnu11* cKO inhibits cell death and delays DNA damage. **(a)** Immunohistochemistry (IHC) for γH2AX (green), CC3 (magenta), and TUNEL (yellow) on sagittal sections of control (left), *Rnu11* cKO (middle), and dKO (right) pallium across E12.5 (A’), E13.5 (A’’), E14.5 (A’’’). Nuclei are marked by DAPI (blue). Scale bars=30µm. **(b)** Quantification of cell death via TUNEL (top) and CC3 (bottom) signal in control, *Rnu11* cKO, and dKO pallium across E12.5, E13.5, and E14.5. Data are presented as mean± s.e.m. Statistical significance was determined by one-way ANOVA, followed by post-hoc Tukey test. n.s.=not significant, *=*P*<0.05; **=*P*<0.01; ***=*P*<0.001. (C) Quantification of DNA damage via γH2AX signal in control, *Rnu11* cKO, and dKO pallium across E12.5, E13.5, and E14.5. Data are presented as mean± s.e.m. Statistical significance was determined by one-way ANOVA, followed by post-hoc Tukey test. n.s.=not significant, *=*P*<0.05; **=*P*<0.01; ***=*P*<0.001.

## Discussion

Unlike mouse models of microcephaly linked to mutations in single genes regulating cell cycle, the microcephaly in *RNU4ATAC*-diseases, including MOPD1, RS, and LWS, are linked to the disruption of the minor spliceosome. As the minor spliceosome is responsible for the removal of minor introns found in 699 human and 650 mouse genes, perturbations will likely have a broad molecular footprint (Olthof et al., 2019). Thus, the underlying the cellular defects that result in microcephaly in such patients proves more difficult to disentangle than most characterized presentations of primary microcephaly. We have previously reported that *Rnu11* cKO pallium has cell cycle defects and cell death caused by misspliced MIGs involved in disparate functions besides the most enriched function of cell cycle regulation (Baumgartner et al., 2018). Importantly, non-MIGs that were upregulated in the *Rnu11* cKO, enriched for only one function being intrinsic apoptotic signaling pathway in response to DNA damage by p53 class mediator (Baumgartner et al., 2018). This finding is consistent with those reported in numerous mouse models of microcephaly, where blocking cell death through genetic ablation of *Trp53* rescued microcephaly (Insolera et al., 2014; Little and Dwyer, 2019). Here we show that ablation of *Trp53* in the *Rnu11* cKO failed to prevent microcephaly (Fig. 1C-D). This finding suggests that microcephaly in the case of minor spliceosome related diseases might not follow the molecular and cellular pathways observed in other microcephalies. Unlike mutations in single genes that alter cell cycle, which account for the majority of cases of genetically linked primary microcephaly, inhibition of the minor spliceosome affected 178 MIGs via minor intron retention (Baumgartner et al., 2018; Jayaraman et al., 2018).Considering this data, it is not surprising that ablation of p53 did not fully rescue in the dKO. The limited rescue we did see, is most likely a consequence of the delay in cell death of NPCs that results in neuron production that is absent in the *Rnu11* cKO mice. Moreover, it is also possible that newly born neurons that might have died in the *Rnu11* cKO, might be spared in the dKO where cell death is blocked.

The similarity in the transcriptome profiles of the dKO to control by PCA suggested that transcriptional changes that separated the cKO were driven by p53 activation (Fig. 2B-D). Thus, the genes downstream of p53 activation such as *Phlda3, Bbc3*, and *Aen* were not differentially expressed in the dKO, which were upregulated in the cKO (Fig. 2F). *Pmaip1* was the only gene that was upregulated in the dKO compared to the control, the TPM values for this gene in both samples is below 1TPM, which we use as threshold of expression (Fig. 2F, Table S1).

### Disruptions of minor intron splicing result in missplicing of MIGs involved in cell cycle

The ablation of p53 shifted the overall gene expression patterns in the dKO to that of the control, yet we did not observe significant rescue of the microcephaly phenotype (Fig. 1). The primary defect of minor spliceosome inhibition resulted in significant elevated minor intron retention, which was elevated at similar levels in the *Rnu11* cKO and dKO compared to the control (Fig. 3A). This was further corroborated by PCA on minor intron retention (%MSI retention), which separated out the *Rnu11* cKO and the dKO from the control (Fig. 3B). In agreement, we found similar AS events in the dKO compared to the control (Fig. 3). Recently we have reported that minor spliceosome inhibition also results in AS across minor introns, which was observed in peripheral blood mononuclear cells of RS, LWS, early-onset cerebellar ataxia (EOCA) patients and *Rnu11* cKO pallium (Olthof et al., 2021). Higher numbers of AS events were observed in the dKO, suggesting that more AS events could build-up when cell death was blocked (Figure 3D). Consistent with our previous reports, mis-spliced MIGs, in both the *Rnu11* cKO and the dKO, highly enrich for cell cycle-related terms (Figure 3F; Figure S2). We find at least 40 shared mis-spliced MIGs that are involved in the regulation of various stages of the cell cycle, underscoring the large footprint of minor spliceosome defect on cell cycle regulation (Figure 3F). This finding can explain why despite blocking cell death, we see minimal rescue of microcephaly in the dKO.

### P53 ablation revelaed that disruptions in mitotic progression initiates cell cycle defects in the cKO

At E12.5 in both the *Rnu11* cKO and dKO pallium, we observe an increase in the percent of RGCs in the prometaphase. However, at this timepoint we already observe a decrease in the number of RGCs in metaphase in the dKO (Fig. 4A&B). This trend continues where decreases in the number of RGCs in telophase is detected a day earlier at E13.5 in the dKO compared to the *Rnu11* cKO (Fig. 4A&B). The early detection of these mitotic shifts in the dKO, can be interpreted as a consequence of the loss of p53, in that RGCs stuck in prometaphase in the *Rnu11* cKO would be subjected to apoptosis (Fig. 6), and suggest that the first cell cycle defect upon minor spliceosome disruption is mitotic defects (Fig. 7B). Surprisingly, at E12.5 we did not observe any significant changes in the percentage of cells in S-phase or in the lengths of S-phase and cell cycle in the dKO (Fig. 5A-B and S2). This suggests that loss of p53 might ameliorate some of the cell cycle defects, or that this observation was due to the absence of cell death, which would normally occur in the *Rnu11* cKO. Consistent with the latter idea, in the *Rnu11* cKO at E13.5 we no longer observe the increase in S-phase and cell cycle length (Fig. 5A&B). However, at this timepoint, we observed a significant increase in the cell cycle length as well at the length of G2-G1 in the dKO (Fig. 5A&B). To explain the delay in this cell cycle length exaggeration, one can invoke the earlier mitotic defects of exaggerated prolongation of prometaphase in the dKO, which may contribute to the calculated increase in G2-G1 length (Fig. 4). Furthermore, at E13.5 we observe no change in the percent of RGCs in S-phase *Rnu11* cKO, which can be explained by accelerated cell death. However, in the dKO at this timepoint we observe a decrease in the percent of RGCs in S-phase and thus no change in S-phase length. Based on the calculation method by Martgyar et al. the cell cycle length was extrapolated to be on average (∼280hrs) in the dKO, which is physiologically not feasible, therefore we called these cells to be stuck in the phases in which they are found (Caviness et al., 1995; Cai et al., 1997; Martynoga et al., 2005). Most likely these cells undergo p53-independent apoptosis at E15.5 (Fig. S3).

**Figure 7.**
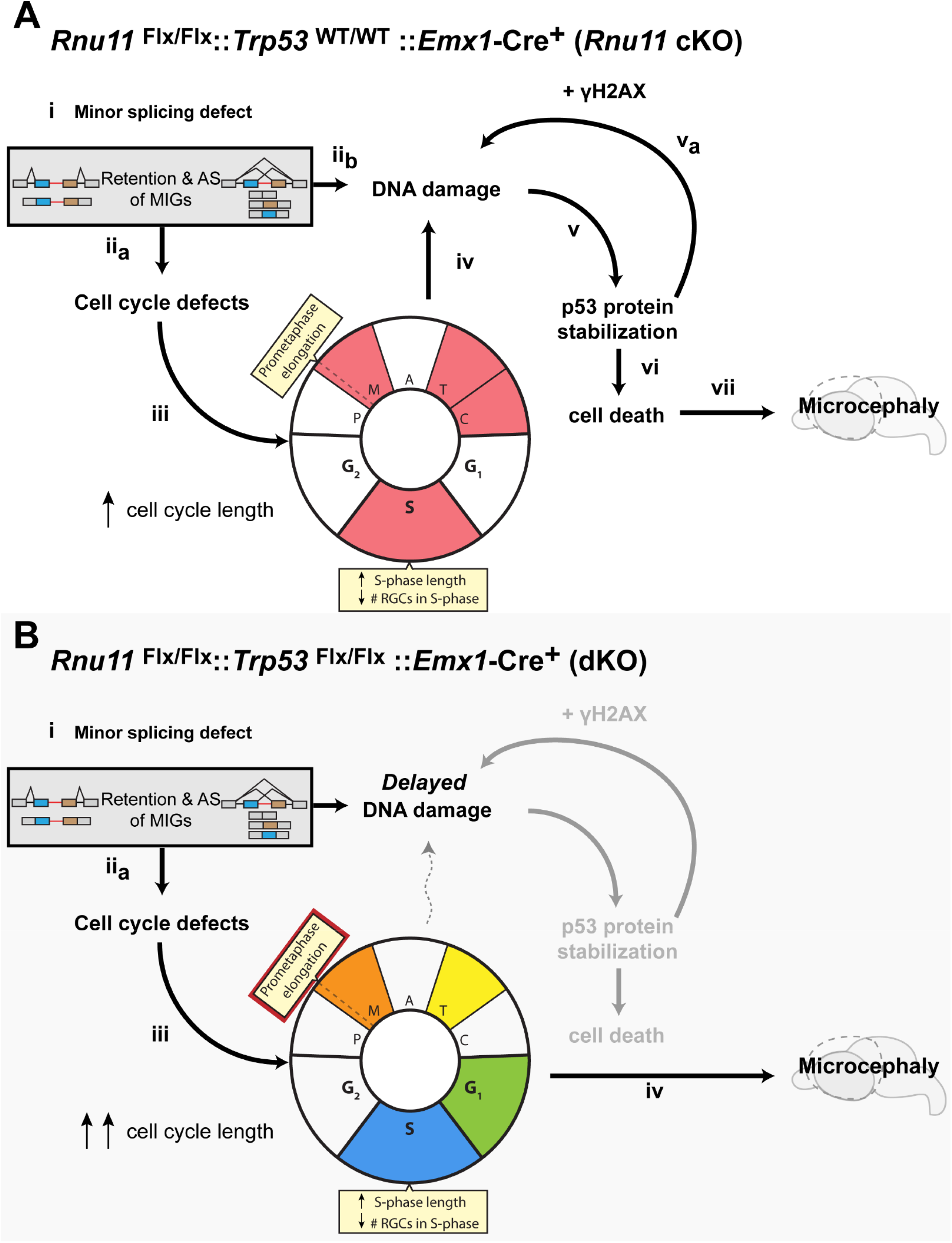
Model of microcephaly progression upon disruption of the minor spliceosome in the context of p53-mediated cell death. **(a)** Summary of microcephaly progression with upon the disruption of the minor spliceosome in the *Rnu11* cKO, with insights from previous work and this study (Baumgartner et al., 2018; Olthof et al., 2021). **(b)** Microcephaly progression upon the disruption of the minor spliceosome in the absence of cell death via *Trp53* ablation in the dKO. Greyed out regions indicate nodes rescued upon genetic ablation of *Trp53*. Color coding of cell cycle defects determine temporal kinetics (red-first to blue-last) of cellular defects that occur as a result of mis-splicing of MIGs.

In short, through knocking out *Trp53* in the *Rnu11* cKO we establish the temporal kinetics of the cell cycle defects leading to microcephaly upon disruptions in minor splicing. Our previously published works support the hypothesis that upon minor splicing defects, as analyzed through minor intron retention and AS around minor introns, occur in cell cycle related genes leading to simultaneous cell cycle defects and DNA damage (Fig. 7A, i, ii_a_, ii_b_). We observed cell cycle defects including elongation of prometaphase, followed by shortening of metaphase and telophase, a reduction in cytokinetic midbodies, and S-phase defects (Fig. 7A, iii). These defects, along with the primary defect of mis-splicing of MIGs involved in DNA damage and repair mechanisms culminate in DNA damage (Figure 7A, ii_b_, iv). This DNA damage then triggers p53 protein stabilization, which further amplifies DNA damage deposition through γH2AX and leads to cell death (Fig. 7A. v, v_a_, vi). Ultimately the cell cycle defect and cell death of RGCs leads to primary microcephaly in the *Rnu11* cKO mice (Fig. 7A, vii).

The current study provides us insight into the more precise temporal kinetics of the cell cycle defects occuring in the *Rnu11* cKO, by blocking cell death. We observe similar primary defects of the mis-splicing of MIGs involved in cell cycle in the dKO (Fig.7B i), which results in cell cycle defects, albeit with shifted kinetics (Fig. 7B, ii_a_). In the dKO, p53 ablation exacerbates the mitotic defect. Here we observe the prometaphase defect a day earlier than in the *Rnu11* cKO, and death at G1 is blocked (Fig. 4 and 7B, iii, grey node). Instead, these RGCs are stuck in G1 or later phases in the cell cycle (Fig. 5 and 7B, iii). Furthermore, we observe increased severity in the S-phase defects in the dKO, including decreased number of RGCs in S-phase and a marked increase in the length of S-phase for those RGCs able to progress to this phase of the cell cycle (Fig. 4 and S2 and 7B, iii). In the dKO, we do not observe significant cell death, but by E14.5 and E15.5 we eventually observe signifcant DNA damage accumulation (Fig. 6C and S3A&B). Ultimately this suggests that microcephaly caused by disruptions in the minor spliceosome are primarily driven by defects in cell cycle (Fig. 7B, iv).

### MIGs affected in the U11-null pallium are individually linked to diseases associated with microcephaly

The core set of 244 MIGs mis-spliced between the *Rnu11* cKO and the dKO enrich for functions including cell cycle and DNA damage response by DAVID analysis (Fig. 3F). This finding suggests that the 244 MIGs underpin microcephaly observed in both *Rnu11* cKO and dKO. Broadly, through generating the *Rnu11* cKO and the dKO mice, we were seeking to understand how perturbations in the minor spliceosome could result in microcephaly pathology. Given that we detected mis-splicing of MIGs in a shared set MIGs and that we observe primary microcephaly in both conditions, we sought to use this core mis-spliced MIG list to inform pathology. To do this, we investigated whether these MIGs harbored mutations that were linked to microcephaly. Indeed, 19 of the 244 MIGs affected in both the *Rnu11* cKO and the dKO themselves lead to diseases in humans characterized by microcephaly, highlighting that these core mis-spliced MIGs may drive microcephaly progression in minor spliceosome disease (Olthof et al., 2020).

### DNA damage, cell death and p53 ablation in the context of disruptions of the minor spliceosome

Ablation of *Trp53*, blocked cell death which most likely allowed us to detect these exaggerated cell cycle defects in the dKO. This is supported by the absence of CC3, TUNEL + cells across E12.5 to E14.5. Surprisingly, we did not observe γH2AX until E14.5 and E15.5 in the dKO (Fig. 6 A&C, Fig. S3). This finding is not consistent with the idea that the accumulation of DNA damage is the trigger that activates p53, which results in apoptosis. Thus, p53 might participate in DNA damage in the context of disruption of the minor spliceosome. This idea is supported by evidence in cancer cells where in a triple knockout (*Rb, p107, p130*) MEFs in which *Trp53* was ablated showed reduction in DNA damage (Benedict et al., 2018). This suggests that in the *Rnu11* cKO, p53 stabilization in response to DNA damage may feedback to further hasten the accumulating DNA damage, however in the dKO, the absence of p53 reduces the speed at which DNA damage accumulates. In all through this study we reveal that microcephaly due to perturbations in the minor spliceosome is primarily driven by cell cycle defects, while shedding light on the interplay between cell cycle defects, DNA damage, activation of p53, and cell death and its relationship to pathology.

## Materials and Methods

### Animal Husbandry

Mouse husbandry and procedures were carried out in accordance with protocols approved by the University of Connecticut Institutional Animal Care and Use Committee, which operates under the guidelines of the U.S. Public Health Service Policy for laboratory animal care. The *Rnu11* conditional knockout mouse used in this study was generated and described by Baumgartner et al. 2018 (Baumgartner et al., 2018). *Emx1*-Cre was bred into the *Rnu11* cKO line to target *Rnu11* for removal in the developing forebrain (Gorski et al., 2002). *Trp53*^*ftm1Br*^ mice were obtained from Jackson Laboratory (JAX stock #008462). For matings intended for embryonic harvests, E0.5 was considered noon the morning a vaginal plug was observed. The experiments described above used male and female *Rnu11*^WT/Flx^::*Trp53*^WT/WT^::*Emx1*-Cre^+^ (control), *Rnu11*^Flx/Flx^::*Trp53*^WT/WT^::*Emx1*-Cre^+^ (*Rnu11* cKO), and *Rnu11*^Flx/Flx^::*Trp53* ^Flx/Flx^ ::*Emx1*-Cre^+^ (dKO) embryos.

### TUNEL Assay

TUNEL was performed on 16 µm sagittal dorsal telencephalon cryosections (E12.5-E14.5) and 16µm coronal embryonic head cryosections (E15.5) using the *in situ* cell death detection kit, TMR Red (Roche Diagnostics, 12156792910), in accordance with the manufacturer’s instructions.

### Immunohistochemistry

For immunohistochemistry (IHC), 16 µm cryosections sagittal dorsal telencephalon cryosections (E12.5-E14.5) or 16µm coronal embryonic head cryosections (E15.5) were used for IHC as described previously. Primary antibodies were diluted to 1:50 (MoBu-1 clone mouse anti-BrdU, Santa Cruz Biotechnology, sc-51514), 1:100 (mouse anti-Aurora-B/AIM1, BD Biosciences, 611082), 1:200 (mouse anti-γH2AX, MilliporeSigma, 05-636; or 1:300 (rabbit anti-cleaved caspase 3, Cell Signaling Technology, 9665S; rabbit anti-p53, Leica Microsystems, p53-CM5p; rabbit anti-Pax6, MilliporeSigma, ab2237).

### Image acquisition and quantification

IHC-processed slides were imaged using a Leica SP2 confocal microscope, utilizing consistent laser intensity, excitation-emission windows, and gain and offset settings across control, *Rnu11* cKO, and dKO sections. For each channel, confocal imaging settings were optimized for fluorescence in the control section for each processed slide. Further processing was performed on IMARIS v8.3.1 (Bitplane) and Adobe Photoshop CS4 (Adobe Systems). As with the confocal settings, image processing was identical among images of control, *Rnu11* cKO, and dKO sections from the same slide. However, for EdU detection experiments performed on E14.5 pallial sections, EdU signal was very low in *Rnu11* cKO and dKO sections, such that confocal settings suitable to the corresponding control did not capture EdU signal in the *Rnu11* cKO and dKO. Therefore, to identify EdU+ cells in the *Rnu11* cKO and dKO pallium, confocal and IMARIS settings were tailored to these genotypes for quantification purposes. Manual quantification was performed in IMARIS as described in Baumgartner et al. 2018 (Baumgartner et al., 2018).

### Cell cycle speed and S-phase speed BrdU, EdU injections

Timed-pregnant dams were injected with BrdU and EdU (100nmol per gram body weight) as described in Martynoga et al., and schematized in Figure 5 (Martynoga et al., 2005). Detection of BrdU was performed through IHC using the MoBu-1 clone mouse anti-BrdU antibody, which does not cross-react with EdU (Santa Cruz Biotechnology, sc-51514; MobU-1(Liboska et al., 2012)). Detection of EdU was performed using the Click-iT EdU Cell Proliferation kit with Alexa Fluor 647 dye (Thermo Fisher Scientific, C10340) in accordance with the manufacturer’s instructions.

### RNA and cDNA preparation

Pallia were dissected from E13.5 control, *Rnu11* cKO, and dKO embryos (n=4 per genotype). Tissue for each embryo was placed in separate aliquots of 100µL TRIzol (Invitrogen, #15596026) and triturated. RNA isolation was then extracted using the DirectZOL RNA MiniPrep kit (Zymo Research, R2050), in accordance with the manufacturer’s instructions. For cDNA preparation for RT-PCR experiments, 300ng of RNA was used for cDNA synthesis as described by Baumgartner et al. 2014 (Baumgartner et al., 2014). Primers used for alternative splicing in Fig. 3E listed in Supplemental Table 2.

### RNAseq

RNA samples were prepared as described in the previous subsection. Library sample preparation and sequencing were executed by the University of Connecticut’s Center for Genome Innovation. Specifically, cDNA library preparation was performed using the Illumina TruSeq Stranded Total RNA Library Sample Prep Kit (RS-122-2201) with RiboZero for ribosomal RNA depletion. Sequencing was performed on the Illumina NextSeq 500 platform. Reads were mapped to the mm10 genome (UCSC genome browser) using Hisat2 (Kim et al., 2015). Isoform and gene expression was determined through IsoEM2, to produce expression values in TPM (Mandric et al., 2017); differential gene expression was then determined using IsoDE2, as previously described (Baumgartner et al., 2018). Minor intron retention and ORF analyses were performed as described in Baumgartner et al. 2018, while alternative splicing around the minor intron was calculated and assessed as described in Olthof et al. 2021 (Olthof et al., 2021). DAVID was employed for functional enrichment analysis of gene sets, with a significance cut-off of Benjamini-Hochberg adjusted *P*-value<0.05 (Huang da et al., 2009). The data reported in this publication have been deposited in NCBI’s Gene Expression Omnibus and are accessible through GEO Series accession number GSE168366.

### Principal component analysis

Principal component analysis (PCA) was performed based on protein coding gene expression (TPM values), retention levels (%MSI_Ret_) using the default settings in ClustVis (Metsalu and Vilo, 2015).

### Nissl staining and neocortical quantifications

P0 brains were cryosectioned (50µm) and collected on 1% gelatin and chrom alum-coated glass slides at 200µm (control) or 150µm (*Rnu11* cKO and dKO) intervals. Slides were rinsed in distilled water for 1 minute, followed by a sequence of ethanol (EtOH) dehydration steps for 2 minutes each (50% EtOH, 70% EtOH, 95% EtOH, 100% EtOH, 100% EtOH, 100% EtOH). This was followed by two 2-minute toluene washes, and then by a sequence of dehydration steps in ethanol, each lasting 2 minutes (100% EtOH, 100% EtOH, 100% EtOH, 100% EtOH, 95% EtOH, 70% EtOH, 50% EtOH). Slides were rinsed in distilled water for 2 minutes, followed by 3 minutes in Cresyl Violet (1% cresyl violet + 0.25% glacial acetic acid), and another 2 minutes water rinse. The first dehydration series was repeated, followed by 3 toluene washes (2 minute, 1 minute, and 1 minute in length, respectively). Slides were mounted with Permount mounting medium (Fisher Scientific, SP15-100) and allowed to harden in the fume hood overnight before being imaged and stitched with Keyence BZ-X710 at 4X.

The medial thickness and mediolateral lengths of the neocortex were measured by ImageJ on images of these coronal P0 brain cryosections. Sections spanning the entirety of the cortex were employed for quantification. For mediolateral neocortical length, the outer edge of the neocortex was measured. Neuroanatomical features and cytoarchitecture, such as the claustrum, rhinal fissure, piriform cortex, amygdala, and shifts in cortical lamination, were used to identify the lateral boundary of the neocortex (Fig. 1D, magenta line)(Franklin, 2008). For medial neocortical thickness measurements, we utilized a line that was parallel to the inter-hemispheric furrow and formed a 90-degree angle with the corpus callosum or, in the absence of the corpus callosum, with the ventricular edge of the neocortex (Fig. 1D, yellow line). For each biological replicate, these values were averaged across the brain.

## Statistical Methods

All statistical tests and results used in this manuscript are outlined in Supplemental Table 3.

## List of abbreviations used

MOPD1: microcephalic osteodysplastic primordial dwarfism type 1
LWS: Lowry-Wood syndrome
RS: Roifman syndrome
snRNA: small nuclear RNA
MIG: minor intron-containing gene
cKO: conditional knockout
dKO: double knockout
NPC: neural progenitor cell
RGC: radial glial cell
AS: alternative splicing

## Notes

### Competing Interest Statement

The authors have declared no competing interest.

